# Segmentation of the Urothelium in Optical Coherence Tomography Images with Dynamic Contrast

**DOI:** 10.1101/2021.02.26.433072

**Authors:** Zhuo Xu, Hui Zhu, Hui Wang

## Abstract

**Significance:** Speckle variations induced by intracellular motion (IM) in the urothelium was observed in optical coherence tomography (OCT) images. It is feasible to use the IM as a dynamic contrast to segment the urothelium with only two sequential OCT images. This new method opens the possibility of tracking the distribution of the urothelial cells to identify the microinvasion of bladder tumors.

**Approach:** With fresh porcine bladder tissue, IM was analyzed by tracking speckle variations using autocorrelation function, then quantified with CONTINE algorism to identify the decorrelation time (DT) of the speckle variations. Variance analysis is conducted to show IM amplitude and distribution in the urothelium. The segmentation of the urothelium is demonstrated using tissue samples with and without significantly stretching.

**Results:** Significant speckle variations induced by IM exists in the urothelium. However, the distribution of the IM is heterogeneous. The DTs are majorly distributed between 1ms to 30ms. Using the IM as a dynamic contrast, the urothelium can be accurately and exclusively segmented, even the layer structure of the bladder is invisible.

**Conclusions:** With fresh porcine bladders, we show that IM can be used as a dynamic contrast to exclusively segment the urothelium. This contrast may provide a new mechanism for OCT to diagnosis the invasion of urothelial cancerous cells for the better staging of bladder cancer.

## 1 Introduction

Intracellular motion (IM) is originated from the motion of molecules and organelles in the cytoplasm of eukaryotic cells. Typically, IM is classified as Brownian motion, superdiffusion, and subdiffusion, which are related to temperature, the viscosity of the surrounding medium, and the density of the cytoskeleton ^1^. IM is essential for the proper functioning of cells. The technology of tracking IM can be divided into two categories. Fluorescence labeling methods are based on fluorescence recovery after photobleaching^2^, fluorescence correlation spectroscopy^3^, and time-resolved fluorescence imaging^4^ to quantify IM, such as intracellular diffusion coefficients or trajectories of labeled particles. Instead of tracking specific molecules, coherent gated methods measure the speckle variation induced by IM. Initially, holographic optical coherence imaging was used to image the IM of tumor spheroids and drugs' responses^5,6^. Later on, with optical coherence tomography (OCT), IM has been used as an endogenous contrast to reveal the cellular and subcellular structures with freshly excised tissue. These cellular and subcellular structures are hardly visible in typical OCT images but have different mobilities, which can be extracted through standard deviation or power frequency analysis ^7,8^. Autocorrelation function analysis and CONTIN algorism also have been employed to quantify IM ^9,10^. The uniqueness of the coherent gated method is that it can detect IM at different depths without requiring fluorescence tagging. Therefore, the imaged objects can stay at a more natural status.

In previous studies, IM has been employed as a contrast to probe cell viability, test drugs, and detect cellular and subcellular structures in tissue. Here, we show that IM can be used to segment the urothelium, a tissue lining the lumen surface of the bladder. Conventionally, the urothelium is considered as an impermeable barrier to separate the urine and the blood. Bladder cancer, the fourth most common cancer among men, is originated from the urothelium^11^. Urothelial cancerous cells are initially confined in the urothelium and may gradually invade the below lamina propria (LP) and the musculus propria (MP). Therefore, tracking the invasion depth of the urothelial cancerous cells is the cornerstone of stratifying patients into different cancer stages, which have a significant impact on treatment strategies. Previous clinical studies show OCT is a promising technology for bladder cancer diagnosis^12–14^. Under OCT, normal bladder tissue after sufficient distention shows a clear boundary between the urothelium and the LP. The boundary is faded or disappeared when the urothelial cancerous cells have invaded the LP. At the same time, typical OCT images based on photon scattering intensity lost the tracking of the urothelial cancerous cells. They cannot identify the invasion depth and patterns of the urothelial cancerous cells anymore. In the last decade, aquaporins, or water channels, have been discovered on the urothelial cells^15–18^. Although the exact function is still not clear, these aquaporins seem related to water transport. With fresh porcine tissue samples, we have directly visualized the dynamic process of water absorption and extraction in the urothelium (manuscript in preparation). After absorbing water, we have observed significant IM in the urothelium. This paper characterizes the IM in the urothelium through autocorrelation analysis of speckle variations and quantifies the distribution of speckle decorrelation time (DT) with CONTINE algorism. Due to the short speckle DT, we show it is feasible to use the IM as a dynamic contrast to exclusively segment the urothelium with only two sequential OCT images.

## 2 Materials and Methods

### 2.1 Optical Coherence Tomography (OCT)

The OCT used in the studies is based on spectral-domain OCT and was designed and built in-house. A Superluminescent Diode (SLD, Inphenix) centered at 840nm with a full width at half maximum (FWHM) bandwidth of 45 nm was used as the broadband light source. The spectrometer in the system consists of a transmission grating at 1200l/mm at 840nm (Wasatch) and a 2048-pixel linear CCD (AVIIVA SM2). The measured axial resolution is ~7.5μm, and the calculated lateral resolution is ~7.9 μm based on the used collimated lens (AC127-019, Thorlabs), focusing lens (AC250-030, Thorlabs), and the fiber mode field diameter (750HP). The system can acquire images at 20K A-scans/s.

### 2.2 Porcine Bladder Tissue

The porcine bladder tissue was acquired from a local slaughterhouse immediately after the animals were sacrificed. The tissue was kept in 4°C Krebs-Henseleit solution before using. The bladder was first flushed with DI water and then cut into smaller pieces (~ 5cm by 5cm). During imaging, the tissue sample was clipped from the four edges and mounted on four translational stages. By moving the translation stages to four directions with the same distance, the tissue sample can be uniformly stretched.

### 2.3 Autocorrelation Analysis

For autocorrelation analysis, we parked the light spot at the same location on a tissue sample and acquired an M-scan image with 80K A-scans at 20K A-scans/s. The DC background was removed by subtracting a prerecorded image by blocking the light in the sample arm.

The 80K A-scans were divided into 20 segments. The intensity AFC *G(τ, z)* at a specific depth z along an A-scan of each segment was calculated as

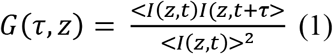

Here, < *I(z, t)* > is intensity by calculating the squared norm of the OCT signal at z.

By assuming random motion of the cell organelles in urothelial cells, the AFC can be fitted with multiple exponential decay functions as

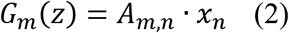

*G_m_(z)* is a vector of the AFC data at z and *x_n_* is the contribution of exponential decay functions with different decorrelation times (DT), γ_*n*_. *A*_*m*,*n*_ is a transfer matrix connecting *G*_*m*_(*z*) and *x*_*n*_ and defined by *A*_*m*,*n*_ = exp(−*t_*m*_/γ_*n*_)*. Solving *x*_*n*_ from equation (2) is an ill-posed problem.

CONTIN algorism addresses this issue by including an additional constraint as a regulation term to seek the minimum value of *D*(*α*) as

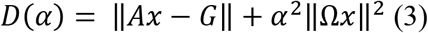

Here, the first term is the residual norm, and the second term represents the cost of the additional constraint, and *α* is the regularizer. An L-curve can be plot as the residual norm versus the additional cost at different values of *α*. The *α* at the corner represents a balance between these two errors and is considered as the optimal value for the best solution^19^. In this study, the additional constraint, Ω, assumes that the size(speckle decorrelation time) distribution of the cellular organelles is continuous and smooth. The ACF data and the optimal *α* is feed into CONTIN algorism to calculate the contribution of different decayed exponential functions.

### 2.4 Variance Analysis

A variance image was calculated from 20 continually recorded frames by scanning the same location. The images were acquired at 20 frames/s with 1000 A-scans across 0.5 mm. The variance of each pixel over 20 frames was calculated and then normalized by the corresponding OCT intensity image.

### 2.5 Segmentation of the urothelium

For segmentation, we acquired 50 images at 20 frames/s, then pick any two sequential images among these 50 images. We adapted the algorism used in optical microangiography to segment the urothelium based on the speckle variation^20^. After removing the DC background, the difference of i^th^ A-scan between the two frames, *D*_*i*_, was calculated as

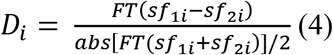

Here, *sf_1i_ and sf_2i_* is the corresponding spectral fringe of an A-scan, and FT is Fourier transform. The difference is normalized to the averaged OCT signal amplitude of the two frames.

## 3 Results

### 3.1 Speckle variation in the urothelium

As shown in Fig.1(a) and (b), we can observe apparent speckle variations in the urothelium with a clear boundary between the urothelium and the LP. However, the degree of the speckle variation is different. Some speckles in the urothelium vary significantly, while others show minimal changes. In contrast, the speckles below the urothelium, such as in the LP, are always stable. IM can observed in video 1.

**Figure 1.**
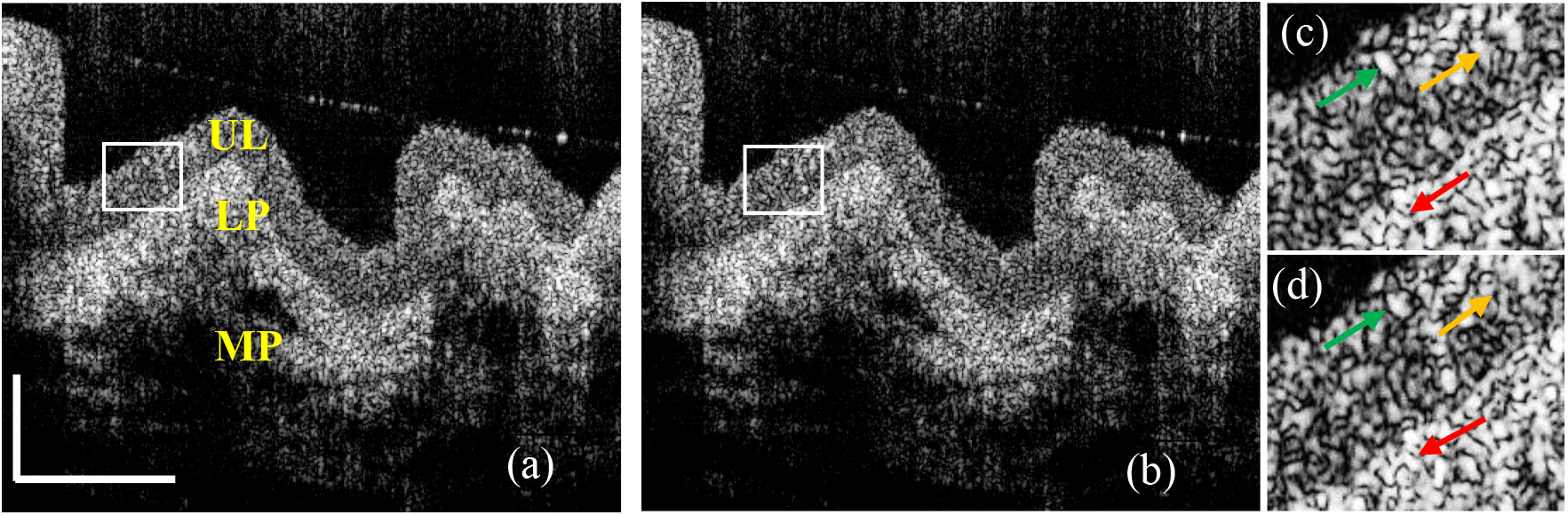
Two OCT images of a fresh porcine bladder tissue sample (a) the first frame; (b) the 11^th^ frame; (c) and (d) are the magnified images cropped from (a) and (b) at the rectangular areas. The yellow arrows point to the speckles with significant changes in the urothelium, while the green arrows point to the speckles with minimal changes. The red arrows point to the speckles, which do not show a noticeable change in the lamina propria. UL: Urothelium; LP: Lamina propria; MP: Muscularis propria (video 1, MP4,1.2MB)

### 3.2 Autocorrelation analysis

By tracking a single speckle with time and calculating the autocorrelation function curve (AFC), we can estimate how fast the molecules or organelles move. Figure 2(a) shows an M-scan image composed of 500 A-scans. The speckle intensity in the urothelium varies with time, while the speckle intensity in the LP or the muscularis propria (MP) below the urothelium is stable. The correlation color map shown in Fig.2(b) is the average of the AFCs of the 20 segments. The pixels of the static tissue show constant AFCs, while the pixels with IM have decayed AFCs. Figure 2(c) shows the AFCs calculated from the M-scan after subtracting the mean intensity value of each pixel and normalized to the autocorrelation value at time delay *τ* = 0. By removing the mean intensity, we kept only the dynamic part of the M-scan. Gradually decayed ACFs with long tails concentrate in the urothelium. As their decay rates are different, the IM at different depths is also different. For static tissue, because the mean intensity values have been removed, the speckle intensity variations mostly come from the random background noise, shown as a sharp peak at τ (delay time) = 0 without significant tails.

**Figure 2.**
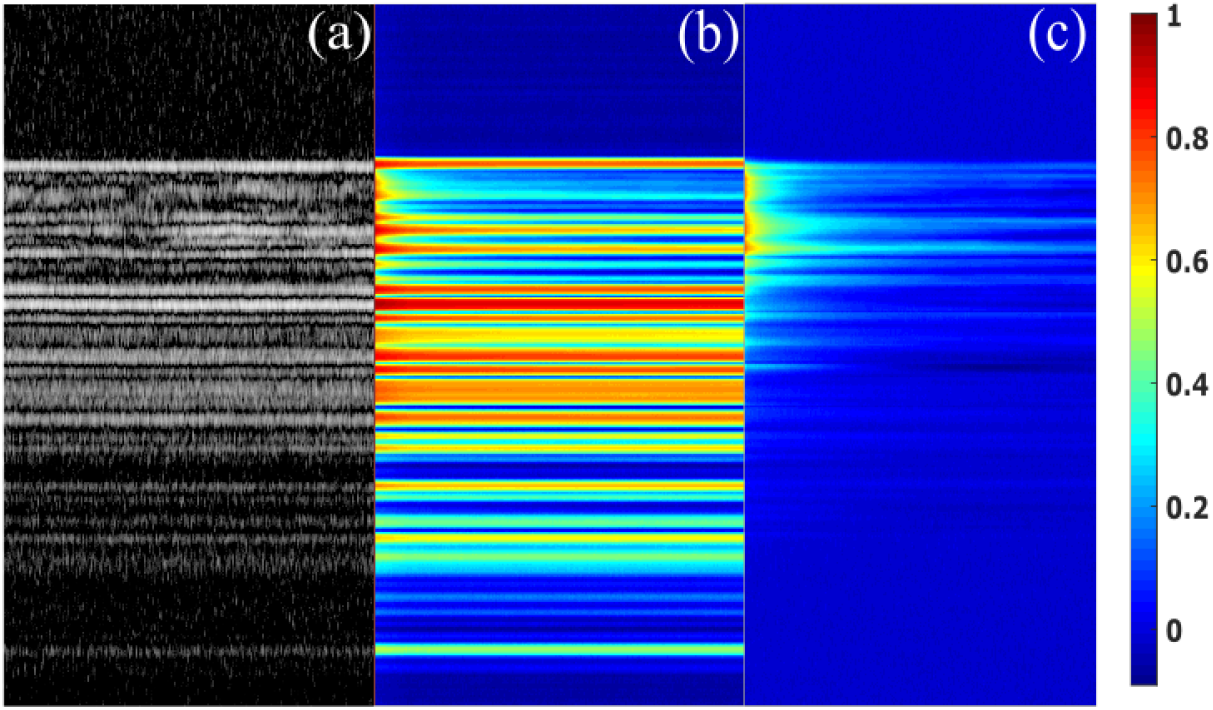
Autocorrelation color maps. (a) An OCT M-scan of a porcine bladder tissue sample; (b) The autocorrelation colormap with the OCT M-scan shown in (a); (c) The autocorrelation colormap after removing the mean value of (a).

CONTIN algorism is used in dynamic light scattering (DLS) to derive the particle sizes of a polydisperse suspension. Here, we employed CONTIN algorism to analyze the ACFs in the urothelium shown in Fig.2(c). Figure 3(a) shows a representative AFC calculated from the speckle intensity variation at a depth in the urothelium. The AFC has two segments. The first segment has a quick drop from 1 to ~0.75 in less than 0.05s, which is shown as the magnified inset in Fig. 3(a). The second segment following the first segment is a gradually decayed tail lasting ~50ms. CONTIN algorism identifies three peaks at 0.165ms, 1.49ms, and 24.6ms, shown in Figure 3(c). These three peaks represent three major DTs, the reciprocals of decay rates of exponential functions. To confirm that the DT peaks have been identified correctly, we recovered the second segment of the AFC with the exponential decay functions having the DTs in Figure 3(c). The AFC and the recovered AFC match very well, as shown in Fig.3(a). In DLS, a decay rate is proportional to a diffusion coefficient. Different DT peaks in Figure 3(b) indicate that the IM captured in the coherence volume of the OCT light beam simultaneously exists three major diffusion coefficients. A longer DT suggests that IM has a slower diffusion. The amplitude of a peak represents the contribution of the DT to the AFC. From Figure 3(b), the speckle intensity variation can be majorly attributed to the IM with DT peaked at 1.49ms and 24.6ms. However, there is a fast IM with DT peaked at 0.165ms. In Fig.3(d), we also show the DT distribution of 100 continuous pixels in the urothelium as a colormap. Clearly, the IM in the urothelium is distributed heterogeneously, while significant DT peaks can be observed in a range between 1ms to 30 ms.

**Figure 3.**
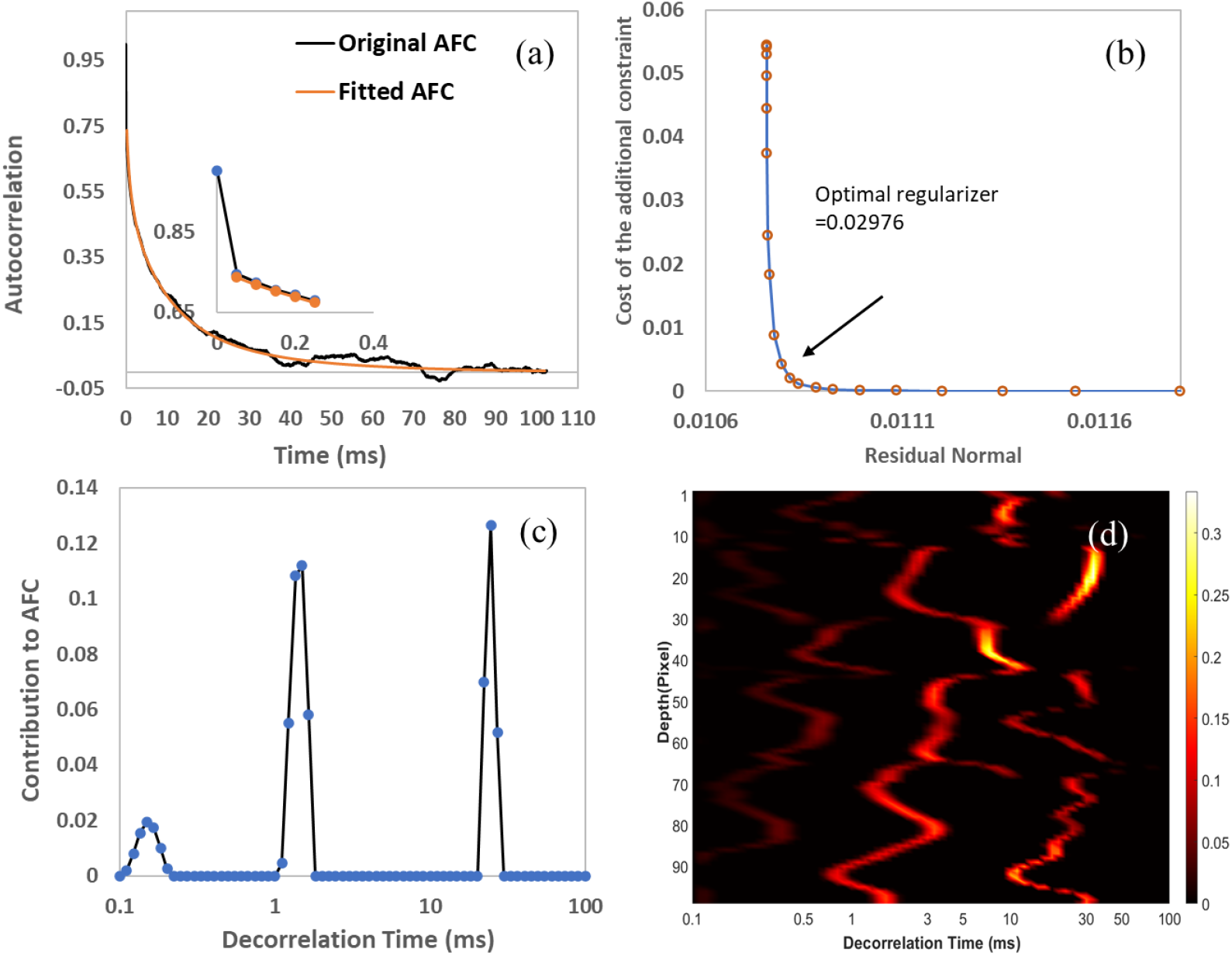
Autocorrelation analysis with CONTIN algorism. (a) A typical autocorrelation function curve (AFC) in the urothelium and the recovered AFC with the decorrelation time peaks identified in (c); (b) L curve used with CONTIN algorism to find the optimal regularizer from the AFC; (c) The decorrelation time derived from the AFC shown in (a) with CONTIN algorism. (d) Decorrelation time color map of 100 pixels in the urothelium calculated with CONTIN algorism; The inset in (a) is the magnified AFC at time delay τ=0.

### 3.3 Variance analysis

We conducted variance analyses to show the amplitude and the distribution of IM in the urothelium. Fig.4(a) is an averaged image from 20 sequentially acquired OCT images. The variance of each pixel is calculated and shown in Fig.4(b). Figure 4(c) shows the fused image of Fig.4(a) and (b). As the boundary between the urothelium and the LP is visible in Fig.4(a), we can find that intense IM matches the urothelium very well from Fig.4(c). However, the IM does not distribute uniformly through the urothelium. By looking closely at the inset image shown in Fig. 4(c), we can find some regions of the urothelium show intense IM, while the other regions only show minor IM similar to those in the LP. Due to the limitation of our OCT system's resolution, we cannot visualize the cellular and subcellular structures, and tell what organelles introduce significant IM with these images.

**Figure 4.**
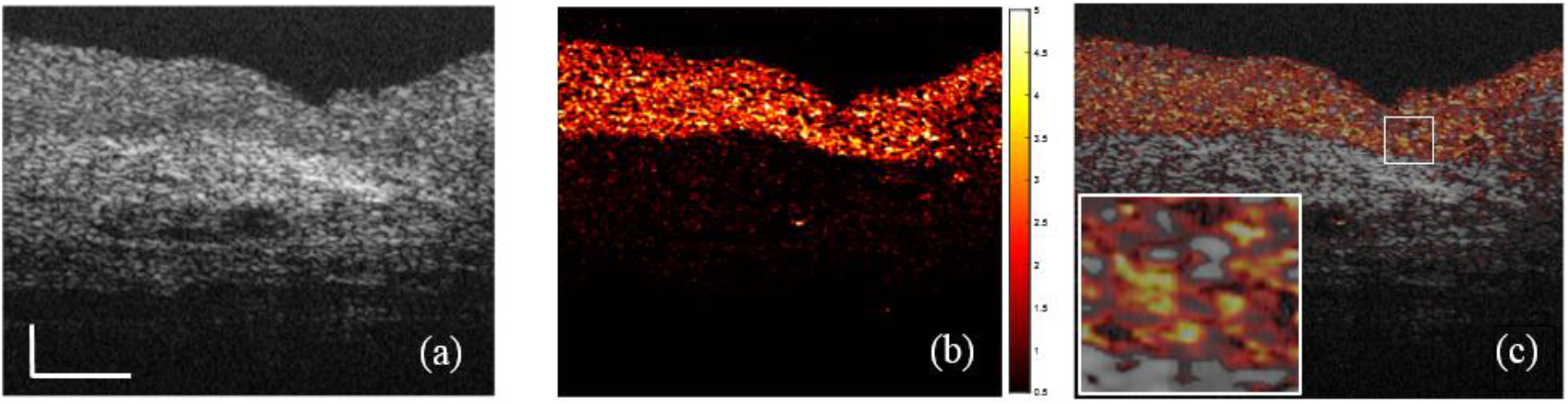
Speckle variance analysis. (a) An OCT image after averaging 20 continuously acquired OCT images at 20frames/s; (b) Normalized speckle variance image of the 20 OCT images; The color map indicates the amplitude of the variance. (c) The fused image of (a) and (b). Scale bar: 0.1mm

### 3.4 Segmentation of the urothelium with intracellular motion

The urothelium is a transitional epithelium, which consists of multiple layers of epithelial cells tightly connected.^21^ The urothelium can be stretched easily to accommodate the volume change of the bladder. When stretched, the urothelial cell tends to become longer, thinner, and flatter. These changes result in a sparser distribution of the organelles in the cells along the bladder surface compared with the non-stretched bladder. Therefore, the light scattering from the urothelium can be significantly reduced after stretching. In contrast, the LP consists of several types of cells, such as fibroblasts, adipocytes, and nerve endings, surrounded by an extracellular matrix (ECM) full of elastic fibers.^22^ When the LP is stretched, the space of the ECM between the cells in the LP tends to be compressed, so the density of the cells in the ECM is increased, leading to enhanced light scattering. Under OCT, a clear boundary can be observed between the urothelium and the LP when the tissue is stretched sufficiently due to different scattering levels. In clinics, the layered structure observed under OCT on the normal bladder is because the bladder has been significantly stretched by filling saline water. However, when a bladder tissue is not stretched or stretched minimally, there is no visible boundary between the urothelium and the LP, then it is difficult to differentiate the urothelium under OCT.

In Figure 4, we have observed that the variance of the IM overlaps with the urothelium very well. In Figure 5, we demonstrate the feasibility of using IM as a dynamic contrast to segment the urothelium with only two sequentially acquired images. Figure 5(a) shows an OCT image of a stretched bladder tissue sample, the boundary between the urothelium and the LP is clear. Figure 5(b) shows the segmented urothelium using the complex difference of two sequential OCT images described in section 2.5. The extracted IM part closely matches the shape and the boundary of the urothelium, which is visible in Figure 5(a). This result gives us the confidence to use IM as a dynamic contrast to extract the urothelium. Figure 5(c) shows an OCT image of a bladder tissue sample with minimal stretch. We cannot observe the boundary between the urothelium and the LP at all due to the minimal stretch. However, using the IM, the urothelium can be extracted exclusively and shown in Figure 5(d).

**Figure 5.**
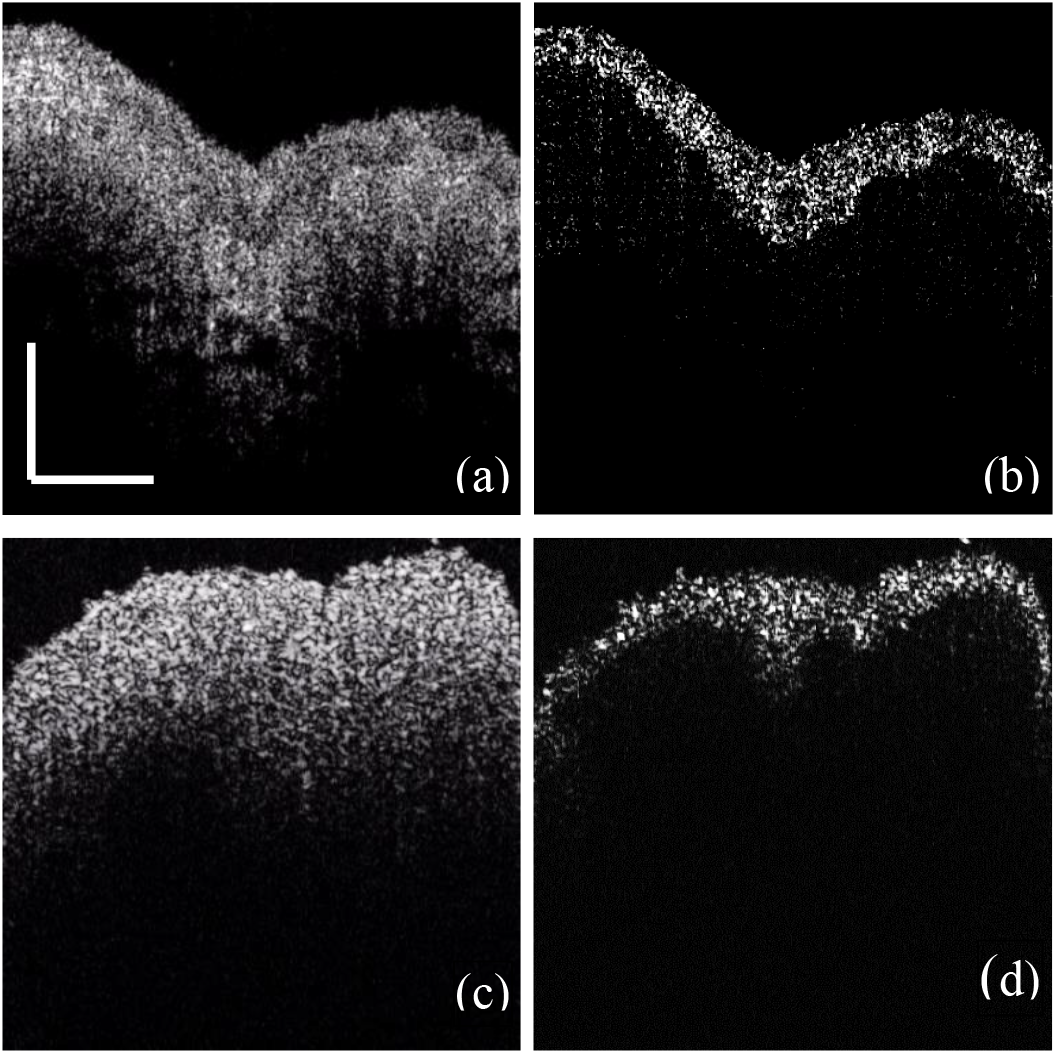
Segmentation of the urothelium with intracellular motion. (a) An OCT image of a fresh porcine bladder tissue sample with gentle stretching; (b) The image reconstructed after the subtraction of the OCT complex signals between the two sequential images like (a); (c) An OCT image of a fresh porcine bladder tissue sample with minimal stretching; (d) The image reconstructed after the subtraction of the OCT complex signals between the two sequential images like (c). (Scale bar:0.5mm)

## 4 Discussion

This paper shows significant speckle variations induced by the IM existing in the urothelium under OCT with freshly excised porcine bladder tissue samples. The quantity analysis through CONTIN algorism based on the AFCs indicates that the major DTs of the IM are distributed between 1ms to 30ms. The amplitude of the IM are not homogeneously and uniformly distributed in the urothelium through variance analysis. Finally, we demonstrated that the urothelium could be segmented with only two sequential OCT images using the IM as a dynamic contrast, even when the boundary between the urothelium and the LP was not visible under OCT.

The urothelium consists of three types of cells, the superficial umbrella cells, the intermediate cells, and the basal cells interfacing with the LP^23^. Conventionally, the urothelium is considered as a passive barrier between the blood and the urine. Transmission electron microscopy reveals that the apical surface of the umbrella cell is covered with plaques, which are interconnected by hinges^24^. Although the function of this plaque-hinge structure is still not clear, they may contribute to the barrier function of the urothelium. Adjacent umbrella cells are tightly bound by tight junctions, which seal the lateral intercellular space to prevent the paracellular flux across the urothelium. However, aquaporins (AQP) have been discovered in the human, rat, and porcine urothelium, challenging the exclusive barrier function of the urothelium and suggesting that water can be transported into the urothelium ^15–18^. Numerous organelles distribute in urothelial cells, such as mitochondria, endoplasmic reticulum, nucleus, glycogen granules, lamellar body filled with lipid droplets, and unique fusiform-shaped vesicles ^23,25^. Due to the limited resolution of the OCT, we cannot directly observe these organelles in images. However, the backscattered photons from the organelles within the coherent volume of an OCT can form speckles shown as irregular patterns. Such speckles are very sensitive to the organelles' motions up to the nanometer scale ^26^. Therefore, the organelles' motions are reflected as the intensity variation of the speckles in OCT images. As long as the surface of the urothelium is covered with a thin layer of water, we can observe significant IM in the urothelium shown in the video1.

Based on the speckle variation, we conducted quantitive analyses of the IM in the urothelium using two methods. The organelles' diffusion rates are varied at different depths of the urothelium. A slow diffusion has a gradually reduced AFC with a long tail, while static tissue has a single-peak AFC at time delay τ = 0 when the mean values were removed. With CONTIN algorism, we can fit the AFCs with exponential decay functions, which are characterized as DT. DT is the duration for dynamical speckles to become uncorrelated. According to DLS^10^, the reciprocal of DT is proportional to the diffusion rate. We calculated the DT distribution of 100 continuous pixels along with the depth of the urothelium. The IM at different depths has three or four DT peaks with major contributions from the peaks between 1ms to 30ms. The multiple DT peaks suggest that several different diffusion patterns exist simultaneously in the coherent volume of an OCT. This observation can be attributed to the different sizes, shapes, and densities of the organelles. **It also implies that as long as two OCT images are acquired with a time interval longer than 30ms, we should observe apparent speckle variations.** However, the AFC analysis cannot directly tell us the motion amplitude of the IM in the urothelium, so we conducted variance analyses. We observed that IM overlapped with the urothelium very well, and the distribution of the IM is heterogeneous. The dynamical regions with large variances are mixed with the static regions showing minimal variances. This observation indicates that not everywhere in the urothelium has observable IM during the 1s imaging time. Besides, intense IM seems mostly located around the middle and the lower part of the urothelium. The top layer of the urothelium is formed by the umbrella cells, which are large polyhedral cells with a size up to 250 μm after stretching ^24^. The umbrella cells have multiple large nucleates (~5-10μm) and are filled with fusiform-shaped vesicles and other organelles. These organelles are mostly located around and below the central part of the umbrella cells ^25^. The lack of dense organelles may explain the less IM in the sub-apical regions of the urothelium.

Segmentation of the urothelium has important clinical significance. Bladder cancer, including papillary tumors and carcinoma in situ (CIS), is originated from the urothelium. OCT has shown potential to differentiate muscle-noninvasive and muscle-invasive bladder cancer base on the visibility of the boundary between the urothelium and the LP, when the bladder is stretched sufficiently ^12,13^. Histologically, this boundary should be the location of the basement membrane^27^, a single layer separating the urothelium and the LP. **The fade or disappearance of the boundary in an OCT image indicates that the urothelial cancerous cells have invaded the LP**. In other words, typical OCT images with scattering as a contrast cannot further stage muscle-invasive bladder tumors, such as differentiating T1a, T1b, and >T2 tumors. Recently, the invasion pattern and depth of the urothelial cancer cells from the basement membrane, termed as microinvasion, has been correlated with the prognosis of bladder cancer ^28,29^. Clearly, it is clinically significant if OCT can exclusively track **urothelial cells no matter the boundary exits or not**.

We have shown that we could segment the urothelium pretty well by calculating each pixel's variance with 20 sequential images. However, this method is not realistic in clinics due to the long image acquisition time. Then, using only two sequential images, we showed that the urothelium could be exclusively extracted using the IM as a dynamic contrast, even when the boundary is not visible. The contrast comes from the phase and intensity differences of the dynamic speckles between two sequential images. Our imaging interval is ~50ms longer than the maximum DT, ~30ms. As we acquired images at 20 frames/s, the method could be adopted in clinics. A high imaging speed can reduce the influence of human body motions on segmentation. We believe that further increasing the imaging speed to above 30 frames/s is also possible. CIS is flat but highly risky bladder cancer ^30,31^. Detection CIS is challenging under white light cystoscopy^32^. Under OCT, CIS shows blur and discontinued basement membrane due to cytologic changes of the urothelial cells, but often confound with inflammatory tissue^13,14,27,30^. Exclusivly tracking of the urothelial cells with the dynamic contrast may allow us to visualize their distribution and then improve the specificity of detecting CIS.

Our study has shown that intense IM exists in the urothelium. The speckle variations induced by the IM can be used as a dynamical contrast to exclusively segment the urothelium no matter the boundary between the urothelium and the LP is visible or not. As we already know that the boundary is faded or disappeared when urothelial cancerous cells start invading the LP and the MP, this capability may open a new route to *in vivo* detect microinvasion in papillary bladder tumors and CIS. However, due to the limited resolution and contrast, we cannot identify the organelles that contribute to the IM. Combining fluorescence imaging with OCT may help us to answer this question in the future. All images in this study were *ex vivo* obtained with fresh porcine tissue as done by other dynamical imaging studies ^6–8^, it should be recognized that the IM may be dominated by the Brownian motion for excised tissue. For *in vivo* studies, super diffusion (active transport), such as motor-driven organelles through microtubules, may play a vital role^33,34^. The more intense *in vivo* IM may allow us to employ OCT with imaging speed at ~100 fps to further avoid potential motion artifacts. Therefore, it is critical to verify the results through *in vivo* studies in the future. Besides, it is essential to conduct comparative studies using normal and cancerous human bladder tissue and correlate OCT images with corresponding histopathological analyses for translating the technology into clinics

In conclusion, we have observed intense IM existing in the urothelium. Quantitively analyses show that the IM with major DT were peaked between 1ms and 30ms and is heterogeneously distributed in the urothelium. Such intense IM can be used as a dynamical contrast to exclusively segment the urothelium with only two sequential OCT images. We believe this new contrast can be used *for in vivo* tracking the invasion of the urothelial cancerous cells to the LP and the MP.

## Disclosure

No conflict of interest is declared.

## Acknowledgments

This research is supported by U.S. Department of Defense (DOD) Medical Research Program Discovery Award (W81XWH1810104)

## Notes

### Competing Interest Statement

The method for bladder cancer diagnosis

